# Consumption of a high energy density diet triggers microbiota dysbiosis, hepatic lipidosis, and microglia activation in the nucleus of the solitary tract in rats

**DOI:** 10.1101/2020.02.21.949354

**Authors:** Dulce M. Minaya, Anna Turlej, Abhinav Joshi, Tamas Nagy, Patricia Di Lorenzo, Andras Hajnal, Krzysztof Czaja

**Affiliations:** Department of Veterinary Biosciences and Diagnostic Imaging, University of Georgia. Athens, GA, 30602; Department of Pathology, University of Georgia, Athens, GA, 30602; Department of Psychology, Binghamton University, Binghamton, NY; Department of Neural and Behavioral Sciences, the Pennsylvania State University, College of Medicine, Hershey, PA

**Keywords:** High energy density diet, high fat diet, microbiome, gut dysbiosis, leptin, cytokines, hepatic steatosis, microglia, vagus

## Abstract

Obesity is a multifactorial chronic inflammatory disease. Consumption of high energy density (ED) diets is associated with hyperphagia, increased body weight and body fat accumulation, and obesity. Our lab has previously shown that short-term (4 weeks) consumption of a high ED diet triggers gut microbiota dysbiosis, gut inflammation, and reorganization of the gut-brain vagal communication. The aim of the current study was to investigate the effect of long-term (6 months) consumption of high ED diet on body composition, gut microbiome, hepatocellular lipidosis, microglia activation in the Nucleus of the Solitary Tract, and the development of systemic inflammation. Male Sprague-Dawley rats were fed a low ED diet (5% fat) for two weeks and then switched to a high ED (45% fat) diet for 26 weeks. Twenty-four hour food intake, body weight, and body composition were measured twice a week. Blood serum and fecal samples were collected at baseline, and 1, 4, 8, and 26 weeks after introduction of the high ED diet. Serum samples were used to measure insulin, leptin, and inflammatory cytokines using Enzyme-linked Immunosorbent Assay. Fecal samples were used for 16S rRNA genome sequencing. High ED diet induced microbiota dysbiosis within a week of introducing the diet. In addition, there was significant microglia activation in the intermediate NTS and marked hepatic lipidosis after four weeks of high ED diet. We further observed changes in the serum cytokine profile after 26 weeks of high ED feeding. These data suggest that microbiota dysbiosis is the first response of the organism to high ED diets and this, in turn, detrimentally affects liver fat accumulation, microglia activation in the brain, and circulating levels of inflammatory markers.

## 1. Introduction

Obesity has been widely recognized as a low-grade, chronic inflammatory disease. However, the causal relationship between inflammation and obesity, and the causal factors behind obesity-dependent inflammation are not well understood. Inflammation is a transient physiological response of an organism to reinstate homeostasis in response to a, typically harmful, stimuli. The inflammatory state that accompanies the metabolic syndrome observed in most obese individuals, however, is not transient.

Consumption of high energy density (ED) diets, including diets high in sugar, and/or in fat have been shown to trigger microbiota dysbiosis, increased body fat accumulation, and development of metabolic syndrome^1, 2^. The vast majority of obesity cases result from an imbalance between energy intake and energy expenditure, with intake surpassing expenditure. The excess energy consumed is taken up and stored as adipose tissue. Once thought to be inert tissue, adipose tissue is now recognized as a metabolically active endocrine organ regulating physiological and pathological processes^3^. Some of the hormones, adipokines, and cytokines, e.g IL-6, TNF-α, released by adipose tissue have pro-inflammatory or anti-inflammatory effects.

Excess energy intake leads to enlargement of adipose tissue depositions. This enlargement occurs through an increase in the number of adipocytes (adipogenesis) or an increase in the size of existing adipocytes (hypertrophy)^4^. In the *db/db* mouse model, it has been shown that in the early stages of obesity development, there is a high number of adipogenic/angiogenic cell clusters. The number of these cell clusters declines as time goes by and there is an increase in the number of crown-like structures, which are hallmarks of local infiltration of macrophages into tissue surrounding dead adipocytes^5^. Using three independent adipocyte-specific anti-inflammatory mouse models, Asterholm *et al*. showed that an acute inflammatory response in adipose tissue is necessary to stimulate adipogenesis as well as proper remodeling and angiogenesis of the extracellular matrix, to allow for healthy adipose tissue expansion^6^. It would appear that the tonic activation of the innate immune system induced by excess energy intake gradually disrupts the homeostatic state of metabolic processes, triggering a chronic inflammatory state.

Studies have shown that in the obese state, the production of pro-inflammatory adipokines like TNFα and IL-6 induce resident macrophages to change their phenotype from surveillance “M2” to pro-inflammatory “M1” as well as trigger recruitment of M1 macrophages^7, 8^. In addition, it has been reported that free fatty acids activate Toll-like receptor (TLR) 4, a pattern recognition receptor expressed on innate immune system cells, in adipose tissue to generate pro-inflammatory signals^9, 10^. Moreover, deletion of TLR5, highly expressed in the intestinal mucosa and thought to help fight against infections, triggered a shift in the species composition of the gut microbiota that is associated with development of metabolic syndrome in mice ^11^. Previous work from our laboratory has shown that four weeks of high ED diet consumption is sufficient to trigger significant activation of microglia, resident macrophages in the brain, in the Nucleus of the Solitary Tract (NTS) – first relay point for satiety signals arising from the gut^12^.

Another obesity comorbidity is Non-alcoholic Fatty Liver Disease (NAFLD). NAFLD is a condition frequently found among people with diabetes (50%) and obesity (76%), and it is almost universal among diabetic people who are morbidly obese^13^. It refers to a wide spectrum of liver damage, ranging from simple steatosis to steatohepatitis, advanced fibrosis, and cirrhosis that occurs in people who drink little to no alcohol. This disease is the most common chronic liver condition in the Western world ^14^. Hepatocytes play a primary role in lipid metabolism. Free fatty acids (FFAs) enter the hepatocyte and the majority of the FFAs are esterified to form triglycerides (TGs). A smaller fraction of FFAs are used to synthesize cholesterol esters or phospholipids, or broken down to produce ketone bodies. TGs are complexed with an apolipoprotein to form lipoproteins and then are exported from the hepatocyte. The apolipoproteins are synthesized by the hepatocyte and this is the rate limiting step in TG export. Interference with any of the steps above can result in accumulation of intracellular lipid (hepatic lipidosis). When consuming a high ED diet, the usual cause of hepatic lipidosis is the increased production of TGs which outpaces apolipoprotein production^15^.

Therefore, the aim of this study was to study the systemic responses to long-term consumption of a high ED diet. We tested the hypotheses that high ED diet consumption induces progressive microbiota dysbiosis and increases body weight, body fat accumulation, and circulating levels of leptin, insulin, and pro-inflammatory cytokines. In addition, we hypothesized that high ED diet consumption induces NAFLD and increases microglia activation in the Nucleus of the Solitary Tract (NTS).

## 2. Methods

### 2.1 Animals

Male Sprague-Dawley rats (n = 15; ∼300g; Envigo, Indianapolis, IN) were housed individually in conventional polycarbonate shoe-box cages in a temperature-controlled vivarium with ad libitum access to standard pellets of rat chow (PicoLab rodent diet 20, product #5053, Fort Worth, TX) and water. Rats were maintained on a 12:12-h light: dark cycle with lights on at 0700-h and allowed to acclimate to laboratory conditions for one week prior to starting experiments. All animal procedures were approved by the University of Georgia Institutional Animal Care and Use Committee and conformed to National Institutes of Health Guidelines for the Care and Use of Laboratory Animals.

### 2.2 Food Intake, body weight, and body composition

Following the acclimation period, rats were maintained on standard chow for an additional two weeks. The animals were then switched to a high energy density (ED) diet (45% calories from fat, Research Diet #D12451, New Brunswick, NJ). Twenty-four hour food intake was measured twice a week for the duration of the study. Briefly, pre-weighed food (∼ 50 g) was provided in standard stainless-steel hoppers. Twenty-four hours later, the amount of food remaining in the food hopper and all spillage was recorded. Body weight and body composition were measured weekly. A Minispec LF 110 BCA Analyzer (Bruker Corp., The Woodlands, TX) was used to measure body composition in minimally restrained, non-anesthetized animals. The Minispec is a body composition analyzer based on time-domain nuclear magnetic resonance technology, which provides absolute masses for fat, lean tissue, and water ^16^. Six rats were sacrificed after being on the high ED diet for four weeks (short-term ED, or STED group). The remaining nine rats were maintained on the high ED diet for a total of 26 weeks (long-term ED, or LTED group). An additional aged-matched, standard chow fed group of rats (n = 9, LF26) served as the end-point controls for the LTED group.

### 2.3 Cytokines, Leptin, and Insulin levels in serum

Blood samples were collected on the last day of standard chow and 4, 8, and 26 weeks after introduction of the high ED diet. Blood was collected from the lateral saphenous vein and allowed to coagulate in the vial. After one hour, the blood was centrifuged at 10,000 rpm for five minutes. The serum was collected and stored at −21°C. A cytokine array (Rat Cytokine ELISA Kit, cat #EA-4006, Signosis Inc., Sunnyvale, CA) was used, according to manufacturer’s instructions, to compare levels of cytokines and chemokines at each time point. Insulin levels were determined using the Rat Insulin ELISA kit (cat #80-INSRT-E01; ALPCO Diagnostics, Inc., Salem, NH) following manufacturer’s instructions.

### 2.4 Microbiome analysis

Fecal samples were collected following the same timeline as for blood samples mentioned above. Bacterial DNA was extracted from feces using a commercial kit following manufacturer’s instructions (Quick-DNA Fecal/Soil Microbe Miniprep Kit, cat #D6010, Zymo research, Irvine, CA). The eluted DNA was sent to the Georgia Genomics and Bioinformatics Core at the University of Georgia for sequencing. High throughput sequencing was performed using Illumina MiSeq paired-end runs. Amplification targeted the V3-V4 region of the 16S ribosomal RNA genes using the following primers: S-D-Bact-041-b-S-17 (5’-CCTACGGGNGGCWGCAG-3’) forward and S-D-Bact-0785-a-A-21 (5’-GACTACHVGGGTATCTAATCC-3’)^17^. Sequences were subsequently trimmed, joined, and quality filtered. To identify Operational Taxonomic Units (OTUs) and to evaluate beta (community diversity divergence between samples) and alpha (microbial diversity within sample) diversities, we used the Quantitative Insights Into Microbial Ecology (QIIME) software package^18^. Linear discriminant analysis to identify taxa with differentiating abundance was conducted using the LDA Effect Size (LEfSe) algorithm^19^. Bacterial abundance was normalized by log-transformation, and statistical analysis and principal component analysis (clustering) were performed using the METAGENassist platform^20^.

### 2.5 Euthanasia and tissue processing

Rats were anesthetized with CO_2_ and transcardially perfused with 0.1 M phosphate-buffered saline (PBS; pH 7.4) followed by 4% paraformaldehyde. Hindbrains and liver were harvested, post-fixed in 4% paraformaldehyde for 2-h, and immersed in 30% sucrose and 0.1% NaN3 (Sigma-Aldrich; pH 7.4) in PBS and stored at 4°C until processing.

### 2.6 Microglia activation

Hindbrains were cryosectioned (Leica CM1950, Leica Biosystems, Wetzlar, Germany) at 20 μm thickness and standard immunofluorescence was used to determine microglia activation in the hindbrain. Hindbrain sections were incubated overnight with a primary antibody against ionized calcium binding adaptor molecule 1 (Iba1, Wako Cat#019-19741, RRDI: AB_839504) followed by Alexa-488 secondary antibody for 2-h to visualize microglia activation as previously described^21^. Sections were mounted in ProLong (Molecular Probes, OR) and examined under a Nikon 80-I fluorescent microscope. The area fraction of Iba1 was analyzed using Nikon Elements AR software as previously described ^22, 23^.

### 2.7 Hepatic Lipidosis

After fixation, liver samples collected from the right median lobe were trimmed, processed, and embedded in paraffin. Paraffin blocks were then cryosectioned at 4 µm thickness and the tissue sections were stained with hematoxylin and eosin (H&E) for intrahepatic lipid content assessment. H&E stained liver sections were examined microscopically by a board-certified veterinary pathologist using an Olympus BX41 upright light microscope.

In addition, samples for Oil-Red-O (ORO) staining were embedded in Optimum Cutting Temperature (OCT) compound (VWR Inc., Atlanta, GA) and were cryosectioned at 7 µm thickness. Tissue sections were mounted on positively charged slides, stained with ORO stain (Polysciences Inc., Warrington, PA), and coverslipped. Histological images were captured using an Olympus DP25 digital camera controlled by Olympus cellSense Standard software at 200x and 400x original magnification (Olympus, Shinjuku, Japan). A semi-quantitative grading scale (normal [0], minimal [1], mild [2], moderate [3], and marked [4]) was used to express the extent of hepatic lipidosis ^24^. In addition, ORO staining was quantified by Nikon Elements AR Software. A threshold was determined to isolate the ORO stain from the background. The threshold settings for 200x were determined as R: 145-255, G: 0-129, B: 2-255, circularity: 0-1, and size: 0-92. Random systematic sampling was completed by drawing five 600 x 600 pixel squares, using the square ROI tool, on the image. The squares were arranged in a manner resembling non-overlapping Olympic rings. The sampling was completed so the squares were randomly assigned to areas uninterrupted by empty space (sinusoids). Binary area fraction was calculated as previously described^23^ and averaged from the five squares. These averages were then compared between groups to analyze changes in hepatic lipidosis.

### 2.8 Statistical analysis

GraphPad Prism 7 (GraphPad Software, Inc.) was used to conduct statistical analyses. Data are expressed as mean ± SD and were analyzed using t-test or ANOVA followed by Holm-Sidak multiple comparisons test as appropriate. Alpha value for statistical significance was set at 0.05.

## 3. Results

### 3.1 High energy density (ED) diet consumption significantly increased body weight and body fat mass

Group means for caloric intake, body weight, and body fat mass were compared using one-way ANOVA to evaluate the effect of short term (STED) and long term (LTED) consumption of a high ED diet (Fig. 1). In the STED group, the animals significantly increased their caloric intake during the first week after introduction of the high ED diet compared to intake of the low ED diet (Ps < 0.0001) [Fig. 1-A]. Caloric intake decreased to intakes of the low ED by week two and remained stable. After two weeks of high ED diet consumption, the animals were significantly heavier than at baseline (P < 0.01) [Fig. 1-B]. After four weeks on the high ED diet, rats were still significantly heavier than after week one on the diet (P = 0.007). Consistently, body fat mass percent significantly increased after only one week on the high ED diet compared to baseline (Ps < 0.05) and remained high in spite of the decrease in caloric intake [Fig.1-C].

**Fig. 1.**
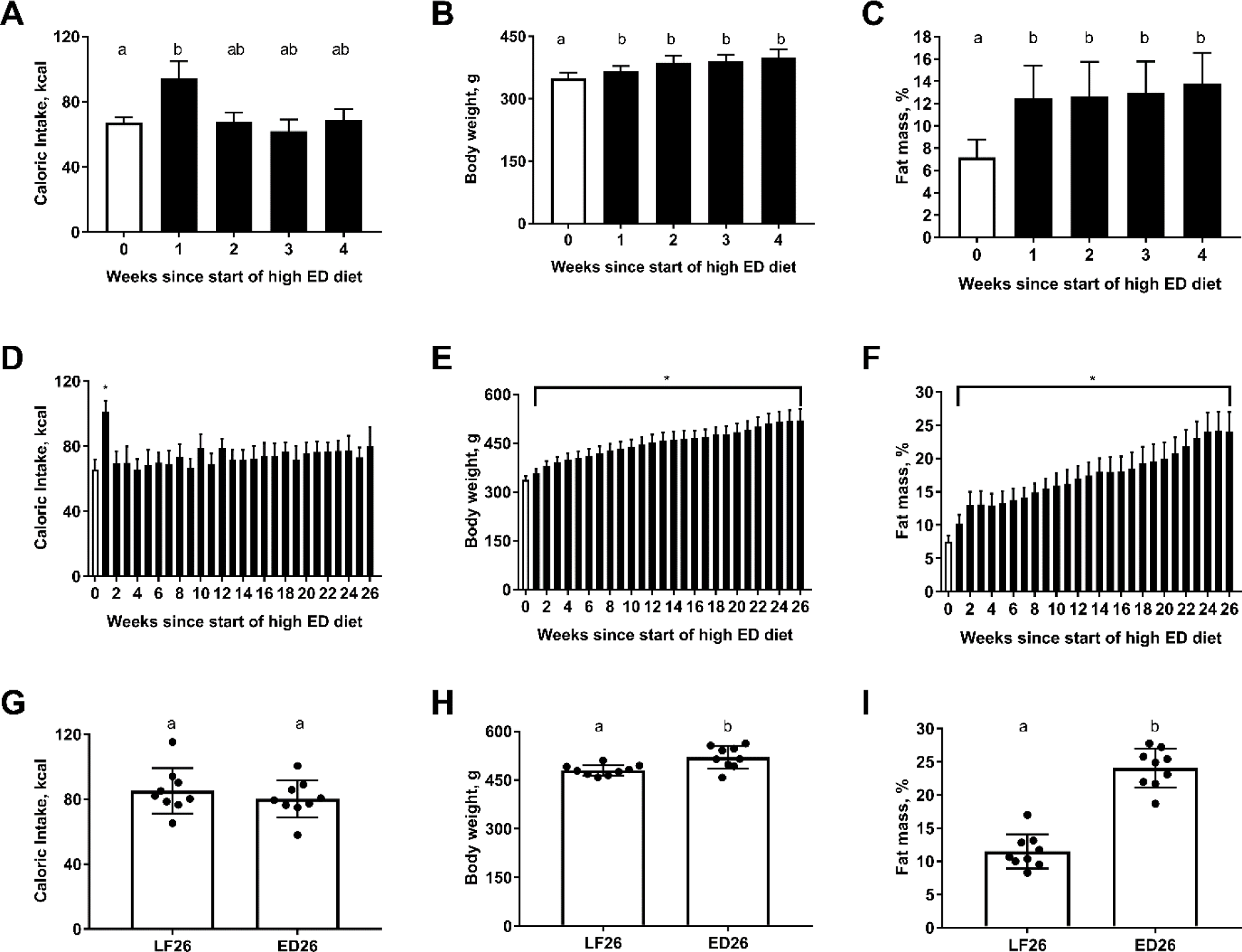
High energy density (ED) diet consumption significantly increased body weight and body fat mass. Shown are mean ± SD kcal consumed (A, D, G), body weight (B, E, H), and body fat mass (C, F, I) for rats fed high ED for four weeks (**STED**; n = 6, top row), rats fed high ED for 26 weeks (**LTED**; n = 9, middle row), and end-point comparison of rats fed low ED vs high ED for 26 weeks (n = 9 per group, bottom row). Animals significantly increased their caloric intake upon introduction of the high ED diet, but caloric intake declined after one week and remained stable for the duration of the study. High ED diet consumption significantly increased body weight and fat mass. Bars denoted with different letters (a, ab, b) differ statistically. Asterisks indicate statistical significance from baseline. P < 0.05.

Similar results were observed in the LTED group. Caloric intake was significantly higher during the first week of high ED diet consumption compared to intake of low ED diet and all subsequent weeks on high ED diet (P < 0.0001) [Fig. 1-D]. Body weight was significantly higher after two weeks of high ED diet compared to baseline (P = 0.037), and the animals continued to gain weight steadily for the remainder of the study [Fig. 1-E]. Body fat percent was significantly higher after two weeks on the high ED diet compared to baseline (P < 0.0001) and fat deposits continued to grow throughout the experiment [Fig. 1-F].

Furthermore, caloric intake of the LTED group at the end of the study (26 weeks of high ED diet) was similar to that of aged-matched, low ED diet controls (LF26) [Fig. 1-G]. However, body weight (P = 0.006) [Fig. 1-H] and body fat (P < 0.0001) [Fig. 1-I] were significantly higher in high ED fed rats compared to low ED diet controls.

### 3.2 Long-term high ED diet consumption significantly changed the cytokine profile in serum

Group means ± SD serum levels of cytokines (OD) and insulin (ng/ml) were compared to investigate the impact of short term and long-term consumption of a high ED diet on [Table 1]. In the STED group, the data were compared using paired t-test. We observed a significant increase in TNFα (P = 0.03) and significant decrease in IL-1α after four weeks on ED diet (ED4) compared to baseline (P < 0.0001). There were no other significant changes observed.

**Table 1.** Cytokine/chemokine optical density values ± SD.

|  | STED |  | LTED |  |  |  |  |
| --- | --- | --- | --- | --- | --- | --- | --- |
|  | LF | ED4 | LF | ED4 | ED8 | ED26 | LF26 |
| **TNFα, OD** | 0.057±0.016 | 0.044±0.011<sup>#</sup> | 0.049±0.014 | 0.044±0.006<sup>*</sup> | 0.038±0.003<sup>#</sup> | 0.057±0.016 | 0.062±0.052 |
| **VEGF, OD** | 0.029±0.006 | 0.026±0.005 | 0.027±0.005 | 0.025±0.004 | 0.019±0.002 | 0.024±0.005 | 0.022±0.011 |
| **FGFβ, OD** | 0.016±0.001 | 0.015±0.001 | 0.014±0.003 | 0.028±0.009<sup>#†</sup> | 0.017±0.007<sup>†</sup> | 0.037±0.007<sup>#</sup> | 0.042±0.042 |
| **IFNγ, OD** | 0.056±0.019 | 0.045±0.009 | 0.055±0.015 | 0.036±0.011 | 0.032±0.005<sup>#</sup> | 0.031±0.004<sup>#</sup> | 0.036±0.031 |
| **Leptin, OD** | 0.049±0.014 | 0.041±0.005 | 0.042±0.002<br>n = 8 | 0.068±0.022<sup>†</sup><br>n = 8 | 0.079±0.071<sup>†</sup><br>n = 8 | 0.472±0.092<sup>#¢</sup><br>n = 8 | 0.089±0.043<br>n = 8 |
| **MCP-1, OD** | 0.027±0.007 | 0.022±0.003 | 0.035±0.007 | 0.08±0.040<sup>#†</sup> | 0.036±0.027<sup>†</sup> | 0.123±0.039<sup>#</sup> | 0.103±0.075 |
| **SCF, OD** | 0.074±0.024 | 0.077±0.038 | 0.064±0.021 | 0.058±0.008<sup>†</sup> | 0.053±0.010<sup>†</sup> | 0.124±0.050<sup>#¢</sup> | 0.063±0.032 |
| **MIP-1α, OD** | 0.019±0.003 | 0.017±0.006 | 0.014±0.002 | 0.074±0.043<sup>#</sup> | 0.043±0.046<sup>†</sup> | 0.108±0.019<sup>#</sup> | 0.089±0.043 |
| **IL-1α, OD** | 0.059±0.005 | 0.027±0.002<sup>#</sup> | 0.054±0.022 | 0.058±0.044 | 0.032±0.006<sup>†</sup> | 0.067±0.026<sup>¢</sup> | 0.024±0.009 |
| **IL-1β, OD** | 0.069±0.017 | 0.056±0.005 | 0.059±0.021 | 0.050±0.009 | 0.043±0.004 | 0.071±0.028 | 0.054±0.032 |
| **IL-5, OD** | 0.059±0.018 | 0.047±0.007 | 0.054±0.016 | 0.032±0.009 | 0.048±0.049 | 0.029±0.007<sup>#</sup> | 0.029±0.019 |
| **IL-6, OD** | 0.021±0.003 | 0.021±0.007 | 0.018±0.003 | 0.022±0.003 | 0.016±0.003 | 0.031±0.011 | 0.024±0.012 |
| **IL-15, OD** | 0.051±0.016 | 0.045±0.010 | 0.046±0.011 | 0.035±0.005 | 0.031±0.005<sup>#†</sup> | 0.046±0.012 | 0.042±0.035 |
| **IP-10, OD** | 0.029±0.004 | 0.032±0.004 | 0.021±0.007 | 0.021±0.004<sup>†</sup> | 0.026±0.003 | 0.029±0.004<sup>#</sup> | 0.027±0.007 |
| **Rantes, OD** | 1.183±0.179 | 0.898±0.169 | 1.191±0.289 | 0.925±0.317<sup>†</sup> | 1.042±0.312<sup>†</sup> | 0.602±0.207<sup>#</sup> | 0.776±0.252 |
| **TGFβ, OD** | 0.279±0.065 | 0.187±0.053 | 0.155±0.012 | 0.091±0.034<sup>#</sup> | 0.198±0.074 | 0.205±0.099<sup>¢</sup> | 0.049±0.009 |
| **Insulin, ng/ml** | 1.223±0.187<br>n = 4 | 1.92±0.346<br>n = 4 | 0.937±0.298 | 1.055±0.185 | 0.888±0.181 | 1.133±0.269 | 1.418±0.486<br>n = 3 |

In the LTED group, longitudinally (data were compared using RM one-way ANOVA), four weeks after introduction of the high ED diet we observed a significant increase in FGFβ, MCP-1, MIP-1a, and TGFβ compared to baseline (Ps < 0.05). After eight weeks of high ED diet (ED8) consumption, serum levels of IFNγ and IL-15 were significantly decreased compared to baseline (Ps < 0.01). Levels of TNFα and IL-6 were significantly lower than after four weeks on the high ED diet (Ps < 0.05). After 26 weeks on the high ED diet (ED26), serum levels of FGFβ, Leptin, SCF, and MCP-1 were significantly higher than at baseline (Ps < 0.05), ED4 (Ps < 0.01), and ED8 (Ps < 0.05). IP-10 levels were significantly higher than at ED4 (P = 0.0094). MIP-1α, IL-15, and IL-1α levels were significantly higher than at ED8 (Ps < 0.05). Serum level of Rantes was significantly lower compared to baseline, ED4, and ED8 (Ps < 0.05). IFNγ, IL-5 levels were significantly lower compared to baseline (Ps < 0.01). Cross-sectional comparison of the ED26 group to low ED diet fed controls using unpaired t-test revealed that the ED26 group had significantly higher levels of leptin, SCF, IL-1α, and TGFβ (Ps < 0.01).

### 3.3 Consumption of a high ED diet triggered progressive dysbiosis of the gut microbiota

To investigate the effect of consuming a high (ED) diet on the gut microbiota, we characterized the microbiome of the STED and LTED groups at several time points throughout the study. In the STED group, we characterized the gut microbiota composition at baseline (LF), one week (ED1), and four weeks (ED4) after introduction of the high ED diet. The rarefraction curve of this analysis indicated > 30000 sequences and > 500 operational taxonomic units per sample [and Supplementary Fig. S1-A]. In the LTED group, we characterized the microbiota composition at four (ED4), eight (ED8), and 26 (ED26) weeks after high ED diet introduction. We also characterized the microbiota of the age-matched, low ED diet control group (LF26). The rarefraction curve of this analysis indicated > 30000 sequences and > 1500 operational taxonomic units per sample [and Supplementary Fig. S1-B].

In the STED group, *Firmicutes* and *Bacteroidetes* were the most abundant phyla and represented >90% of the bacteria identified. The Shannon index, used to characterize species diversity in a community, revealed that there was a significant decrease in bacteria diversity after one week of high ED diet consumption compared to baseline (P = 0.0146) [Fig. 2-A]. There were no significant changes observed in bacterial diversity after four weeks of high ED diet consumption. Principal Component Analysis was used to visualize how different the microbiota were in each sample and to represent the differences as distance [Fig. 2-B]. At baseline (LF), all animals clustered together. One week after high ED diet introduction, the animals clustered away from their baseline (LF) profile. After four weeks of high ED diet consumption, all animals clustered close to the profile at ED1 and further away from their baseline (LF) profile.

**Fig. 2.**
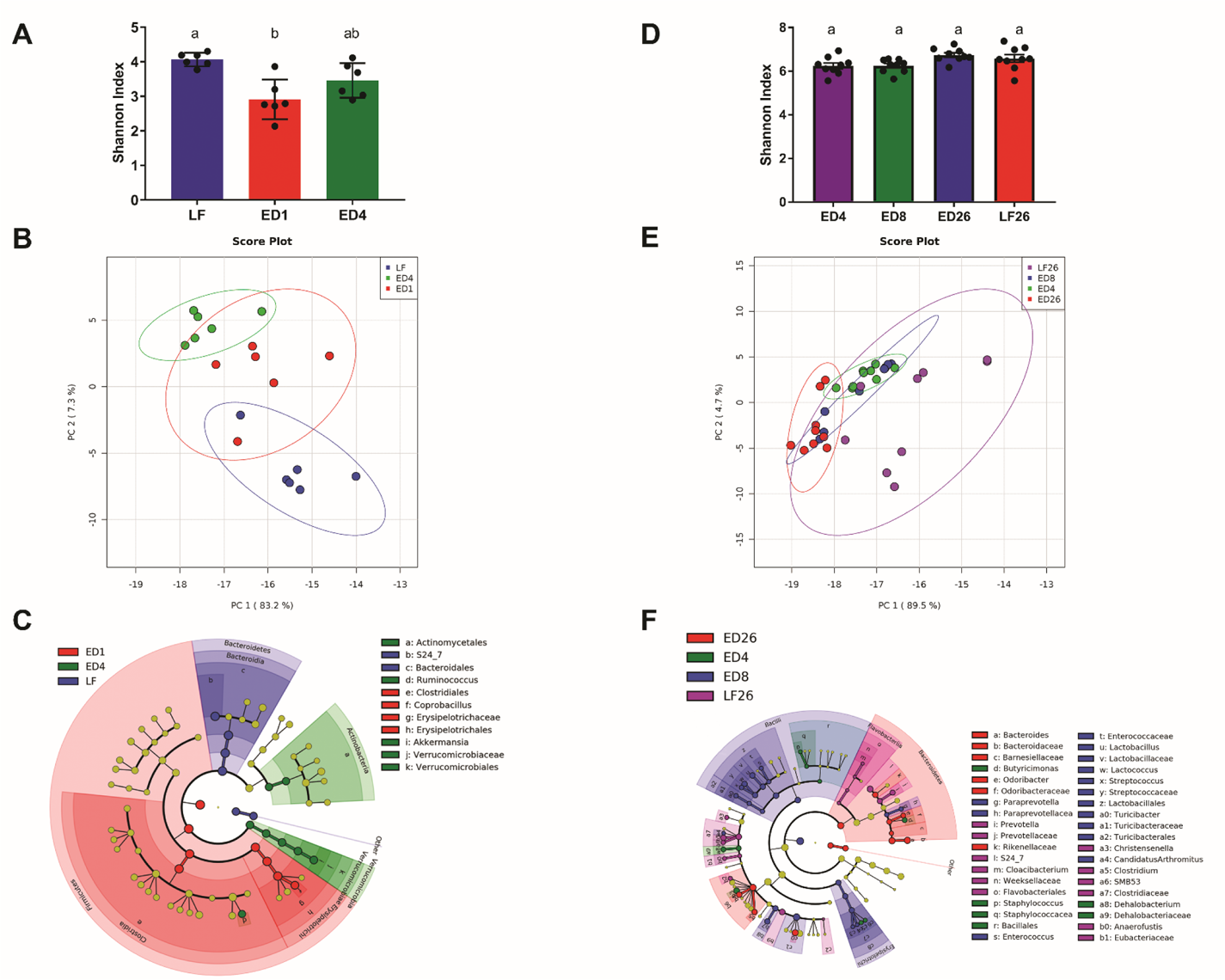
Consumption of a high energy density (ED) diet triggered progressive remodeling of the gut microbiota. The left (A, B, C) and right (D, E, F) columns represent data from the **STED** (n = 6) and **LTED** (n = 9) groups, respectively. **A, D)** Shannon index shown as mean ± SD for each group and time point. Bacterial diversity was significantly decreased after consuming the high ED diet for one week (ED1) compared to baseline (LF). There were no other significant changes observed. **B, E)** Principal Coordinate Analysis showing microbiota from all time points. In the STED group (B), the microbiota of all rats clustered together at baseline (LF). One week after introduction of the high ED diet (ED1), the microbiota clustered together and away from their baseline profile. At week 4 (ED4), the microbiota clustered together and further away from their baseline profile. In the LTED group (E), the microbiota of all rats fed the high ED diet clustered together, independent of time point, and away from the microbiota of rats fed a low ED diet (LF26). C, F) Cladogram produced from LDA scores (see Supplementary Fig. S2 for LDA scores). In the STED group (C), at baseline (LF), the microbiota is characterized by abundant members of the *Bacteroidetes* order *Bacteroidales.* One week after high ED diet introduction (ED1), the microbiota was characterized by abundant members of the *Firmicutes* order *Erysipelotrichales.* After four weeks of high ED diet (ED4), the microbiota was characterized by abundant members of the *Actinobacteria* order *Actinomycetales* and *Verrucomicrobia* order *Verrucomicrobiales.* In the LTED group (F), at ED4 the microbiota was characterized by abundant members of *Bacteroidetes* order *Bacteroidales* and *Firmicutes* orders *Bacillales* and *Clostridiales.* Eight weeks after high ED diet introduction (ED8), the microbiota was characterized by abundant members of the *Bacteroidetes* order *Bacteroidales* and *Firmicutes* orders *Lactobacillales, Turicibacterales,* and *Clostridiales.* After 26 weeks of high ED diet (ED26), the microbiota was characterized by abundant members of *Bacteroidetes* orders *Bacteroidales*. The microbiota of the low ED diet control group (LF26) was characterized by abundant members of *Bacteroidetes* orders *Bacteroidales* and *Flavobacteriales,* and *Firmicutes* order *Clostridiales*.

Figure 2-C [and Supplementary Fig. S2] represents the microbiota composition of the STED group at each time point. At baseline (LF), the microbiota is characterized by abundant members of the *Bacteroidetes* order *Bacteroidales.* One week after high ED diet introduction (ED1), the microbiota was characterized by abundant members of the *Firmicutes* order *Erysipelotrichales.* After four weeks of high ED diet, the microbiota was characterized by abundant members of the *Actinobacteria* order *Actinomycetales* and *Verrucomicrobia* order *Verrucomicrobiales*.

Microbiota composition was changed within a week of introduction of high ED diet [Fig. 3-A]. High ED diet consumption significantly increased the abundance of *Firmicutes* (LF 27% vs ED1 86% and ED4 69%, Ps < 0.0001) and decreased the abundance of *Bacteroidetes* (LF 67% vs ED1 10% and ED4 16%, Ps < 0.0001). There was also a significant increase in abundance of *Verrucomicrobia* after four weeks of high ED diet (LF 1% and ED1 0.8% vs ED4 8%, P = 0.04). At the level of family, high ED diet consumption significantly increased the abundance of members of *Erysipelotrichaceae* (LF 3.8% vs ED1 60% and ED4 41%, Ps < 0.0001) of the phylum *Firmicutes.* Members of the family *S24-7* of the phylum *Bacteroidetes* were significantly depleted by high ED diet consumption (LF 69% vs ED1 12% and ED4 17%, Ps < 0.0001). In addition, one week of high ED diet consumption significantly increased the *Firmicutes-to-Bacteroidetes* ratio, (LF 0.4 vs ED1 8.7, P = 0.0010). There was no statistically significant difference after four weeks on the high ED diet, however, the ratio was still higher at ED4 (4.3) than at baseline (LF) [Fig. 3-B].

**Fig. 3.**
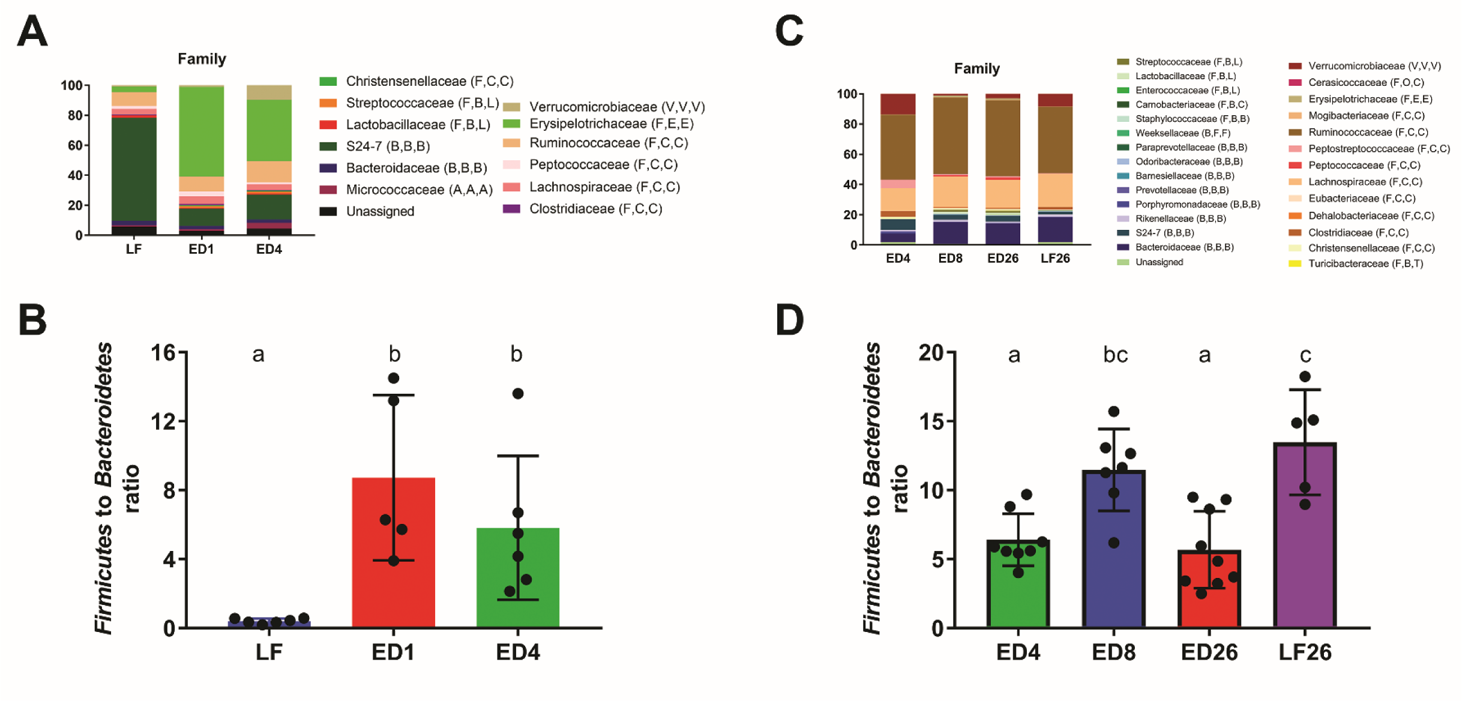
Microbial composition of rats fed a high energy dense diet for 4 weeks (STED, n = 6) or 26 weeks (LTED, n = 9) and rats fed a low energy dense diet for 26 weeks (LF26, n = 9). All phylogenetic levels present with abundance > 1% are represented. **A, C)** relative abundances of phyla at the family level in the STED group and in the LTED. **B, D)** Ratio of Firmucutes to Bacteroidetes in the STED and LTED group. In the STED group, high ED diet consumption significantly increased the abundance of members of *Erysipelotrichaceae* (LF 3.8% vs ED1 60% and ED4 41%, Ps < 0.0001) of the phylum *Firmicutes.* Members of the family *S24-7* of the phylum *Bacteroidetes* were significantly depleted by high ED diet consumption (A). In addition, the ratio of *Firmicutes* to *Bacteroidetes* was significantly higher at one and four weeks of high ED diet compared to baseline (B). In the LTED group, compared to low ED diet fed rats, high ED diet fed rats had significantly higher abundance of members of *Bacteroidaceae* and *Ruminococcaceae,* and significantly lower abundance of members of *Peptostreptococcaceae* and *Verrucomicrobiaceae.*(C). In high ED fed rats, the *Firmicutes-to-Bacteroidetes* ratio was significantly higher after eight weeks compared to after four and 26 weeks (ED8 11.5 vs ED4 6.4 and ED26 5.7, Ps < 0.01). Compared to low ED controls (LF26), high ED fed rats had a significantly lower *Firmicutes-to-Bacteroidetes* ratio at four and 26 weeks (D). In the legend, following the name of each family, higher taxonomic classifications are indicated by letters in parentheses. Phylum: A, Actinobacteria; B, Bacteroidetes; F, Firmicutes; V, Verrucomicrobia. Class: A, Actinobacteria; B, Bacilli if preceded by F and Bacteroidia if preceded by B; C, Clostridia; E, Erysipelotrichia; F, Flavobacteria; V, Verrucomicrobiae; O, Opitutae. Order: A, Actinomycetales; B, Bacteroidales; C, Clostridiales if preceded by C and Cerasicoccales if preceded by O; E, Erysipelotrichiales; F, Flavobacteriales; L, Lactobacillales; T, Turibacterales; V, Verrucomicrobiales. Bars denoted with different letters (a, b, c) differ significantly, P <0.05. Data are means ± SD.

Similarly, in the long-term group (LTED + LF26), *Firmicutes* and *Bacteroidetes* were the most abundant phyla and represented >90% of the bacteria identified. The Shannon index showed no significant difference in bacterial diversity [Fig. 2-D]. Principal Component Analysis showed that all animals fed the high ED diet clustered together and away from animals fed low ED diet (LF26) [Fig. 2-E].

Figure 2-F [and Supplementary Fig. S3] represents the microbiota composition of the LTED group at each time point. At ED4, the microbiota was characterized by abundant members of *Bacteroidetes* order *Bacteroidales* and *Firmicutes* orders *Bacillales* and *Clostridiales.* Eight weeks after high ED diet introduction (ED8), the microbiota was characterized by abundant members of the *Bacteroidetes* order *Bacteroidales* and *Firmicutes* orders *Lactobacillales, Turicibacterales,* and *Clostridiales.* After 26 weeks of high ED diet (ED26), the microbiota was characterized by abundant members of *Bacteroidetes* orders *Bacteroidales*. The microbiota of the low ED diet control group (LF26) was characterized by abundant members of *Bacteroidetes* orders *Bacteroidales* and *Flavobacteriales,* and *Firmicutes* order *Clostridiales*.

At the level of family [Fig. 3-C], four weeks (ED4) and eight weeks (ED8) after high ED diet introduction there was a significantly higher abundance of *Ruminococcaceae* compared to after 26 weeks (ED26, P < 0.01). Four weeks (ED4) and eight weeks (ED8) after high ED diet introduction, there was a significantly lower abundance of *Verrucomicrobiaceae* compared to after 26 weeks (ED26, P < 0.01). High ED diet fed rats had significantly higher abundance of members of *Bacteroidaceae* (LF26 0.8% vs ED4 14%, ED8 14%, and ED26 16%, Ps < 0.0001) and *Ruminococcaceae* (LF26 0.8% vs ED4 14% and ED8 14%, Ps < 0.0001) compared to low ED fed rats. In addition, high ED diet fed rats had significantly lower abundance of members of *Peptostreptococcaceae* (LF26 5% vs ED4 0.6%, ED8 0.9%, and ED26 0.5%, Ps < 0.05) and *Verrucomicrobiaceae* (LF26 14% vs ED4 1.2%, ED8 3%, and ED26 9%, Ps < 0.01) compared to low ED fed rats. In high ED fed rats, the *Firmicutes-to-Bacteroidetes* ratio was significantly higher at eight weeks (ED8) compared to four (ED4) and 26 (ED26) weeks (ED8 11.5 vs ED4 6.4 and ED26 5.7, Ps < 0.01). Compared to low ED controls (LF26), high ED fed rats had a significantly lower *Firmicutes-to-Bacteroidetes* ratio at four weeks (ED4) and 26 weeks (ED26) (LF26 13% vs ED4 6% and ED26 6%, Ps < 0.01) [Fig. 3-D].

### 3.4 Consumption of a high ED diet significantly increased microglia activation in the NTS

We hypothesized that high ED diet consumption triggers microglia activation in the NTS. Results of immunostaining against Iba-1 were compared using one-way ANOVA and revealed that compared to low ED controls (LF26W), rats fed a high ED diet for four weeks (ED4, P = 0.0005) and 26 weeks (ED26, P < 0.0001) had significantly higher binary area fraction of fluorescent staining against Iba-1 [Fig. 4]. In addition, the ED26 group had significantly higher binary area fraction of fluorescent staining than the ED4 group (P = 0.0213).

**Fig. 4.**
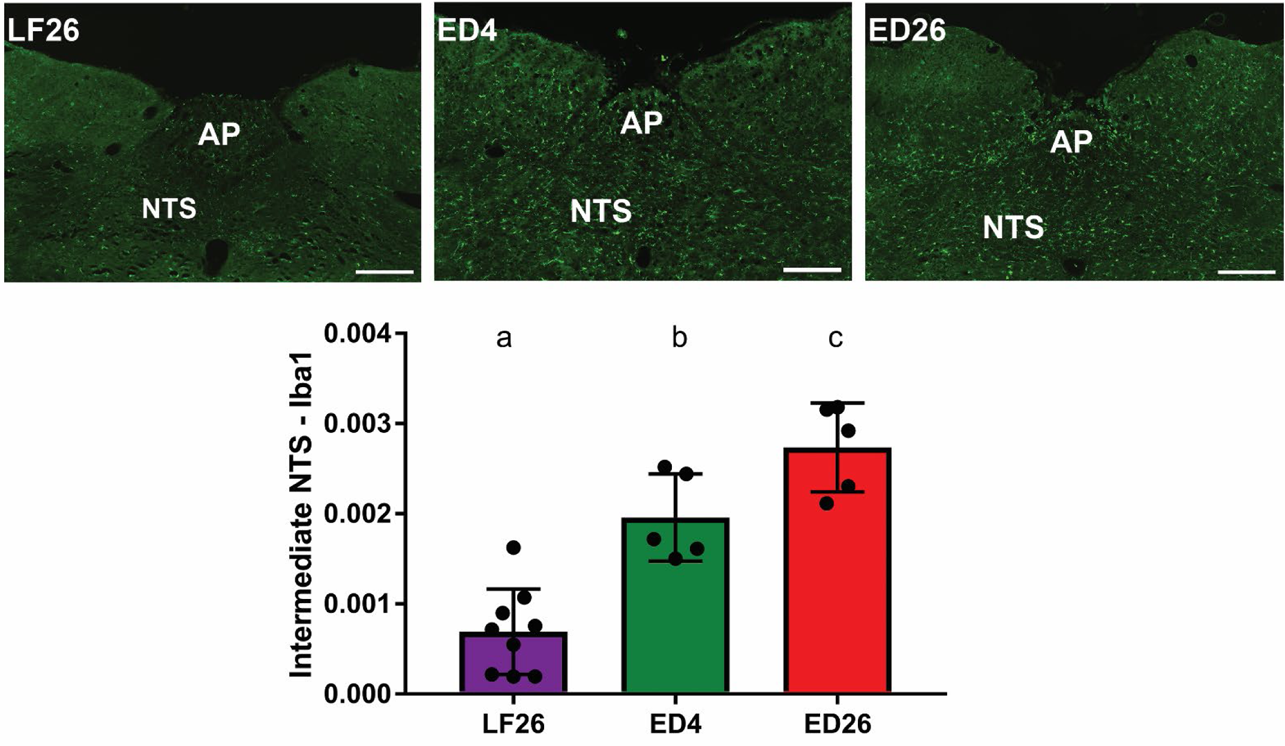
Consumption of a high ED diet significantly increased microglia activation in the intermediate NTS. Representative sections of intermediate NTS of animals fed a low ED diet for 26 weeks (LF26, n = 9), a high ED diet for four weeks (ED4, n = 5), and a high ED diet for 26 weeks (ED26, n = 5) are shown. Binary analysis of the area fraction of Iba1 immunoreactivity showed that animals fed a high ED for four and 26 weeks exhibited significantly more microglia activation than low ED fed controls. In addition, microglia activation after 26 weeks of high ED diet was significantly higher than after four weeks. Graphs represent mean ± SD Iba1 intensity. Bars denoted with different letters (a, b, c) differ significantly (P < 0.05). NTS = Nucleus Tractus Solitarius; AP = Area Postrema. Scale bar = 200 μm.

### 3.5 High ED diet consumption induced hepatic lipidosis

A semi-quantitative method of scoring histological images using Kleiner’s scoring system was completed on ORO and H&E stained hepatic sections with the comparative results shown in Fig. 5. We tested the hypothesis that consumption of a high ED diet leads to increased accumulation of fat in liver tissue. In H&E-stained sections, intracellular lipid is removed during processing and paraffin embedding. Therefore, round, clear, and distinct vacuoles remain in the cytoplasm of hepatocytes. In ORO-stained sections, lipid is retained during embedding and appears as intensely red granules or globules in the cytoplasm of hepatocytes. H&E staining revealed an increase in distinct vacuoles in rats fed the high ED diet (ED4 and ED26) compared to low ED controls (LF26) [Fig. 5, top row]. Similarly, ORO staining showed that high ED fed animals exhibited more intensely red granules (ED4 and ED26) than low ED controls (LF26) [Fig. 5, middle row]. The quantitative scoring of hepatocellular lipidosis confirmed that while animals fed low ED diet (LF26) did not exhibit signs of hepatocellular lipidosis, hepatocellular lipidosis was apparent within four weeks of introducing the high ED diet (ED4). There was no significant difference in hepatocellular lipidosis between the ED4 and ED26 groups [Fig. 5, bottom row table].

**Fig. 5.**
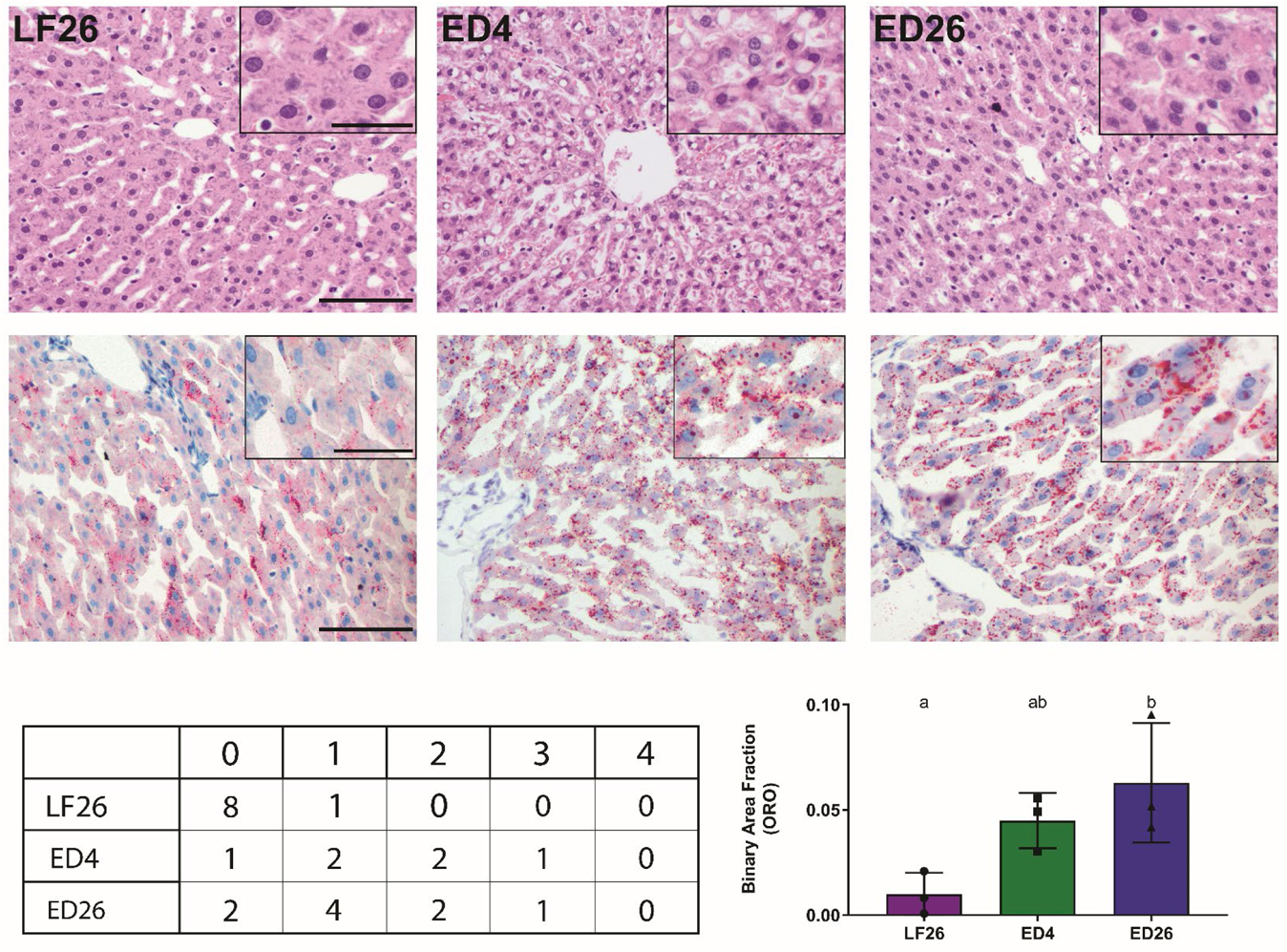
High ED diet intake induces hepatic lipidosis. Representative histopathological images of hematoxylin and eosin stained (top row) and oil-red-o stained (bottom row) hepatic tissue from rats fed a low ED diet (left column), rats fed a high ED for 4 weeks (middle column), and rats fed a high ED for 26 weeks (right column), n = 3 for each group. H&E staining revealed an increase in distinct vacuoles in rats fed the high ED diet (ED4 and ED26) compared to low ED controls (LF26) (top row). Similarly, ORO staining showed that high ED fed animals exhibited more intensely red granules (ED4 and ED26) than low ED controls (LF26) (bottom row). The quantitative scoring (Table) confirmed that hepatocellular lipidosis is apparent after consuming a high ED diet for four weeks. We did not observe significant differences in the extent of hepatocellular lipidosis between the STED (ED4) and LTED (ED26) groups. Low ED fed rats did not show signs of hepatocellular lipidosis. Graph represents mean ± SD binary analysis of the area fraction of ORO staining, which further confirms our results. Bars denoted with different letters (a, b, c) differ significantly (P < 0.05). Scale bar for images is 100 µm. Scale bar of insert is 80 µm.

We further analyzed the intensity of ORO staining in tissue sections that were scored by the veterinary pathologist using a computer software (see methods for details) and results were analyzed using one-way ANOVA. The data revealed that there is a progressive increase in the surface area covered by lipids with regular chow fed animals showing very little staining (BAF = 0.0099 ± 0.01) compared to animals fed the ED diet for four weeks (BAF = 0.0449 ± 0.01) and those fed the ED diet for 26 weeks (BAF = 0.0629 ± 0.02). Animals fed the ED diet for 26 weeks (ED26) had significantly more ORO staining compared to regular chow fed animals (LF26) [Fig.5, bottom row graph].

## 4. Discussion

It is widely recognized that diet is a key factor in short-term and long-term composition, diversity, dynamics, and microbiota-driven host metabolism^25^. In this study, we sought to characterize the long-term effects of consuming a high energy density (ED) diet on the gut microbiome, serum profile of inflammation, neural inflammation, and development of non-alcoholic fatty liver disease (NAFLD). Results showed that high ED diet consumption induced rapid changes in the gut microbiome, triggered inflammation in the NTS as evidenced by increased microglia activation, induced NAFLD, and changed the serum cytokine profile in rats.

Consistent with prior reports ^23, 26–28^, our data showed that upon introduction of a high ED diet, rats significantly increased their caloric intake compared to when fed a low ED diet. However, by the second week of high ED diet consumption the animals had adjusted their caloric intake. This adjustment of caloric intake by rats following acclimation to a high ED diet has been previously reported ^23, 29^. In addition, at the 26 week time point, there was no significant difference in caloric intake between rats fed high ED diet and those fed low ED diet. Concurrently, the rats fed high ED diet exhibited a significant increase in body weight and fat mass that became more prominent over time. Rats fed high ED diet had significantly higher final body weight and fat mass compared to low ED fed animals, despite similar caloric intake. This was previously reported in a study by Lomba *et al*., which showed that rats fed a high fat diet restricted to the amount of calories consumed by a low fat diet fed group gained significantly more body weight and white adipose tissue than the low fat diet fed group^30^. Given that the initial increase in calories consumed by high ED diet fed rats was transient, this phenomenon indicates that high ED diets detrimentally affect body weight and fat mass accumulation independent of caloric intake.

High ED diet consumption induced dynamic fluctuations in the gut microbiota. Consistent with prior reports from our laboratory ^26^, bacterial diversity was significantly decreased within one week of introducing the high ED diet. This immediate response was ephemeral, similar to the transient increase in caloric intake observed during the first week of high ED diet consumption, as we did not detect statistically significant differences in bacterial diversity after four, eight, and 26 weeks of high ED diet consumption. In concert, we observed a similar fluctuation pattern in bacterial abundance. At the one-week mark after introduction of the high ED diet, all animals clustered together and away from their baseline profile. After four weeks of high ED diet consumption, all animals clustered together and away from their baseline and one-week mark profile. Thereafter, we did not observe further fluctuations in bacterial abundance. After four, eight, and 26 weeks of high ED diet, all animals clustered together and away from the profile of animals fed low ED diet.

We found that high ED diet consumption led to a rapid increase in members of the family *Erysipelotrichaceae* belonging to the *Firmicutes* phylum. These are obligate anaerobes have been previously associated with consumption of high ED diets, increased adiposity and inflammation ^31–33^. High ED diet consumption also depleted members of the family *S24-7* belonging to the *Bacteroidetes* phylum and members of the family *Verrucomicrobiaceae* of the *Verrucomicrobia* phylum. The family *S24-7* is associated with gut health as they are primarily involved in the fermentation of dietary fibers to produce short-chain fatty acids (SCFAs) and have been previously shown to be depleted by consumption of high fat diets ^26, 34–36^. Members of the family *Verrucomicrobiaceae* have also been shown to produce the SCFA propionate^35^. By the end of four weeks on the high ED diet, we see a blooming of the family *Ruminococcaceae* that persists for the duration of the study. Members of this family are also SCFAs producers and generally associated with gut health^35^.

In this study, we aimed to characterize the systemic pattern of leptin, insulin, and pro- and anti-inflammatory cytokines in rats fed a high ED diet. We observed a significant increase in serum leptin after consumption of a high ED diet for 26 weeks. We did not observe significant changes at the earlier time points tested. Consistent with prior reports, we did not observe changes in serum levels of insulin with high ED diet consumption at any of the time points tested^37–39^. The effect of high ED diet consumption on insulin levels appears to be strain-specific. Woods *et al*. reported a significant increase in plasma insulin levels in Long-Evans male rats fed a high fat diet for 70 days^40^. Similar results were reported in WNIN rats after consumption of a high fat diet for 13 weeks ^39^ and Wistar rats after 18 weeks ^41^.

The cytokines showing major differences were Fibroblast Growth Factor (FGFβ), Stem Cell factor (SCF), Interferon γ (IFNγ), Interferon γ induced protein (IP-10), Regulated on activation, normal T cell expressed and secreted (Rantes), Monocyte Chemoattractant Protein (MCP-1), Macrophage Inflammatory Protein (MIP-1α), Interleukin 1α (IL-1α), and Interleukin 5 (IL-5). Chemokines, or chemoattractant cytokines, are produced by a host of cell types and play a significant role in inflammatory processes as they help regulate the traffic of immune cells to specific sites. Our results revealed a significant increase in MCP-1 and MIP-1α after 26 weeks of high ED diet consumption. A similar study by Muralidhar *et al*. reported no change in MIP-1α and a non-statistically significant increase in MCP-1 after 13 weeks of high fat feeding^39^. The length of time of high fat/high ED diet consumption likely underlies the differences observed between the two studies, however, there is an observable trend toward higher levels with high fat/ high ED diet consumption. In addition, we observed a significant decrease in Rantes, consistent with a prior report by Fenton *et al*. in mice fed a high fat diet for 10 weeks^42^.

SCF serves as a ligand molecule for the receptor tyrosine kinase c-Kit. Activation of c-Kit by SCF has been shown to be involved in cell migration and survival ^43^. Our results showed a significant increase in SCF levels after 26 weeks of high ED diet consumption compared to baseline. High ED fed rats also have significantly higher levels of SFC than aged-matched, low ED fed rats.

FGFβ is an endocrine hormone produced by the liver, which has been shown to promote gluconeogenesis, ketogenesis, and lipid oxidation during fasting periods. However, during the fed state, FGFβ is thought to enhance insulin-mediated glucose uptake ^44^. Our results showed a significant increase in serum levels of FGFβ after 26 weeks of high ED diet consumption compared to baseline. However, when compared to aged-matched, low ED fed rats, we observed no significant differences. It is thus possible that the changes in FGFβ levels is related to the aging process. To our knowledge, there are no other studies reporting serum FGFβ levels in rats with high ED feeding or in regards to aging.

IFNγ is an important cytokine for innate and adaptive immune response against viral infections. Interferon gamma induced protein 10 (IP10) is a chemokine produced by monocytes, endothelial cells, and fibroblasts in response to IFNγ that acts as a chemoattractant for a hosts of immune cells^45^. Our results revealed a slight decrease in IFNγ and increase in IP10 after 26 weeks of high ED diet consumption compared to baseline. However, there was no significant difference in the level of these cytokines between age-matched high ED fed (ED26W) and low ED fed (LF26) rats. Fenton *et al*. reported a slight, but none significant increase in male mice fed a 35% fat diet for 13 weeks^42^. In Sprague-Dawley rats, Muralidhar *et al*. showed that consumption of a high fat diet for 13 weeks did not affect plasma levels of IP10^46^. It is possible that the longitudinal change observed in this study are a result of the aging process and not necessarily triggered by high ED diet consumption. It has been previously reported that in humans subjects older than 50 years there is a decrease in IFNγ production from mononuclear cells^47^.

Interleukin 1α (IL-1α) is generally known as an epidermal pro-inflammatory cytokine^48^. It has also been shown that IL-1α knockout mice gained less body fat and did not develop glucose intolerance when fed a high fat diet for 16 weeks^49^. Consistent with prior reports^50^, our data showed that rats fed a high ED diet for 26 weeks have significantly higher levels of IL-1α than low ED fed rats. Interleukin 5 (IL-5) is produced by T helper cells and is a key factor in the activation of eosinophils during allergic reactions^51^. Consistent with a prior study^42^, our results showed a significant decrease in serum levels of IL-5 after 26 weeks of high ED diet compared to baseline.

In general, our results revealed longitudinal changes in the serum cytokine profile of animals fed a high ED diet long-term. In addition, when we compared high ED diet fed rats at 26 weeks (ED26) to age-matched, low ED diet fed rats, rats fed high ED diet had significantly higher levels of Leptin, SCF, IL-1α, and TGFβ.

Our research group has conducted several studies to characterize the effect of high ED diet consumption on inflammation in the brain. Vagal afferents carry information from the gut to the NTS. Reports from our laboratory and others have shown that consumption of a high ED diet triggers microglia activation in the nodose glanglia, NTS, and hypothalamus ^27, 52, 53^. Results from this study showed that high ED diet consumption induced an inflammatory response reflected by microglia activation in the intermediate NTS after four weeks. These data further suggest that length of exposure to the high ED diet exacerbates this response since microglia activation after 26 weeks of high ED diet was significantly higher than after four weeks.

Our results demonstrated that consumption of a high ED diet led to a significant increase in intracellular lipid accumulation in the liver, also known as hepatic steatosis. These data are in concert with prior studies in rodents, mice and rats, which report development of hepatic steatosis after consumption of a 40% fat diet for 16 weeks^54^. De Rudder et al., also reported that mice developed hepatic steatosis after only four weeks of consuming a 60% fat diet^55^. Previous studies have shown a link between microbiota dysbiosis and hepatic steatosis^56, 57^. The majority of the blood supply to the liver comes from the intestines through the portal vein^58^. Thus, an increase in gut microbes that produce toxic/inflammatory byproducts increases the gut-derived bacterial products entering the liver^59^. Our data revealed an increase in abundance of members of the family *Erysipelotrichaceae* and a study by Spencer et al. showed that levels of these bacteria are directly associated with changes in liver fat in female human subjects^60^. In addition, we saw an increase in SCFAs producers, namely members of the family *Ruminococcaceae.* The SCFAs acetate, propionate, and butyrate have been previously shown to inhibit lipid accumulation in the liver and improve hepatic function in rodents^61–63^. Given that the abundance of *Ruminococcaceae* was increased after 26 weeks of high ED diet consumption, it is possible that the presence of these bacteria and their byproducts contributed to prevent the progression of hepatic steatosis as there was no difference in the degree of steatosis between the four weeks (ED4) and the 26 weeks group (ED26).

In conclusion, we have shown that long-term consumption of a high ED diet leads to increased adiposity, gut dysbiosis, hepatic steatosis, inflammation in the NTS as revealed by increased microglia activation, and increased systemic levels of inflammatory markers. Our results suggests that gut dysbiosis starts immediately upon introduction of a high ED diet. Next, as the liver is overloaded with increased accumulation of excess fat consumed and toxic bacterial byproducts, hepatic steatosis develops. At the same time, endotoxins produced by the resident gut microbiome damage vagal afferents, which in turn triggers microglia activation in the NTS. Lastly, chemokines, cytokines, and other inflammatory molecules are released from their production site (e.g. adipose tissue) into the systemic circulation. These responses are highly dynamic and play a significant role in the development of obesity and its related comorbidities. In future work, is it necessary to investigate therapeutic interventions to prevent or ameliorate the development of microbiota dysbiosis as this seems to be one of the first and most detrimental consequences of consuming high ED diets.

---

**Supplementary Fig. S1.**
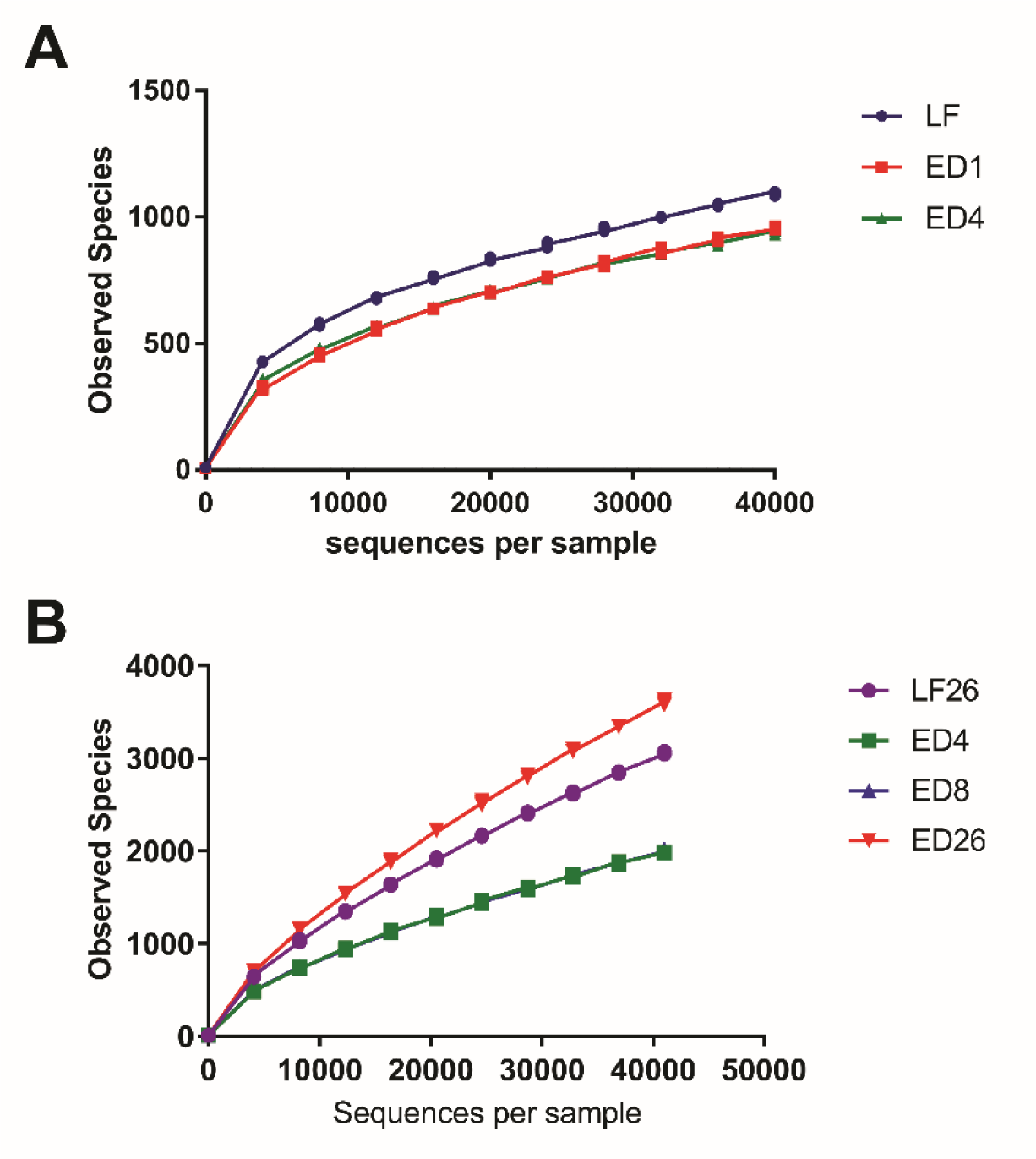
Rarefraction curves by diet group and experimental time point. Data are shown as mean for rats fed a high energy dense diet for 4 weeks (A, STED) or 26 weeks (B, LTED).

**Supplementary Fig. S2.**
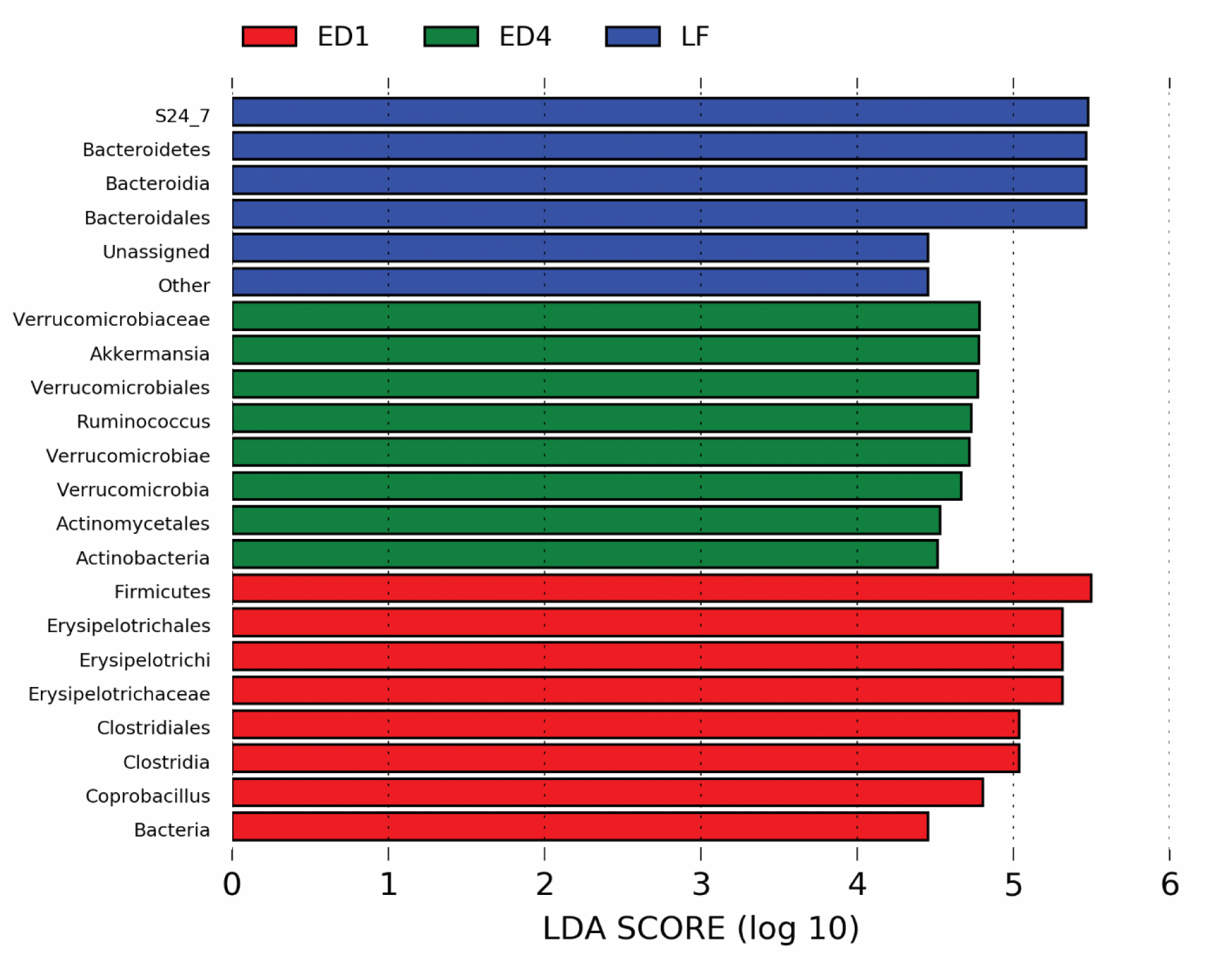
LDA scores used for generation of cladogram (Fig. 3C). Colors designate time point: Blue: LF/baseline, Red: ED1, one week after introduction of high ED diet, Green: ED4, four weeks after introduction of high ED diet.

**Supplementary Fig. S3.**
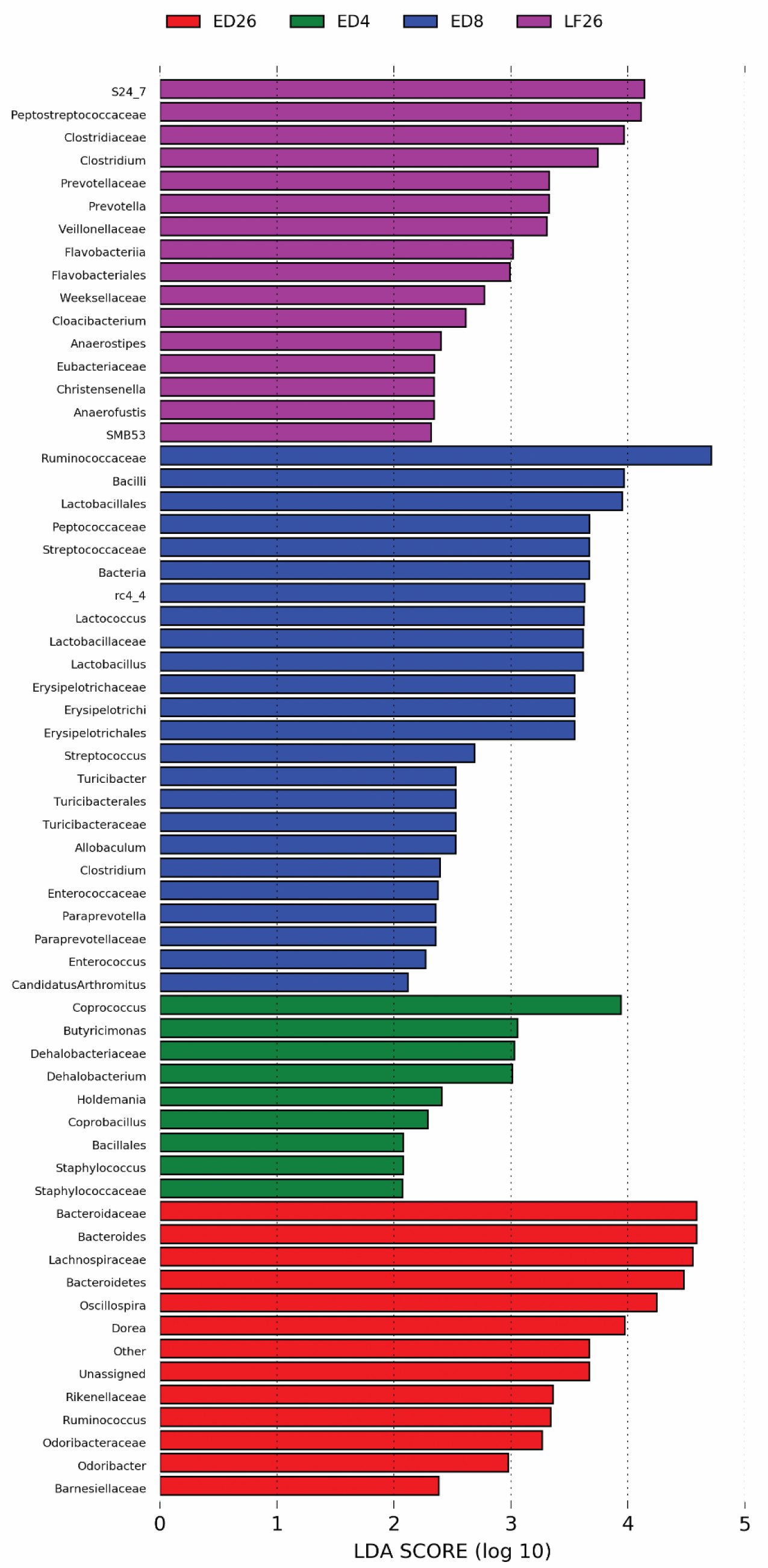
LDA scores used for generation of cladogram (Fig. 3F). Colors designate time point: Purple: LF26, after 26 weeks of low ED diet, Green: ED4, four weeks after introduction of high ED diet. Blue: ED8, four weeks after introduction of high ED diet. Red: ED26, 26 weeks after introduction of high ED diet,

